# Energetics but not development is impacted in coral embryos exposed to ocean acidification

**DOI:** 10.1101/2021.07.19.452948

**Authors:** EE Chille, EL Strand, F Scucchia, M Neder, V Schmidt, MO Sherman, T Mass, HM Putnam

## Abstract

In light of the chronic stress and mass mortality reef-building corals face under climate change, it is critical to understand the processes essential to reef persistence and replenishment, including coral reproduction and development. Here we quantify gene expression and size sensitivity to ocean acidification across a set of developmental stages in the rice coral, *Montipora capitata*. Embryos and swimming larvae were exposed to pH treatments 7.8 (Ambient), 7.6 (Low) and 7.3 (Xlow) from fertilization to 9 days post-fertilization. Embryo and larval volume, and stage-specific gene expression were compared between treatments to determine the effects of acidified seawater on early development. While there was no measurable size differentiation between fertilized eggs and prawn chips exposed to pH 7.8, 7.6, and 7.3, early gastrula and planula raised in reduced pH treatments were significantly smaller than those raised in ambient seawater, suggesting an energetic cost to developing under low pH. However, no differentially expressed genes emerged until 9 days post-fertilization. Notably, gene expression patterns of larvae developing at pH 7.8 and pH 7.3 were more similar than those developing at pH 7.6. Larvae from pH 7.6 showed upregulation of genes involved in cell division, regulation of transcription, lipid metabolism, and oxidative stress in comparison to the other two treatments. While low pH appears to increase energetic demands and trigger oxidative stress, the developmental process is robust to this at a molecular level, with swimming larval stage reached in all pH treatments.

**Summary statement:** This developmental time series tracks the physiological and transcriptomic outcomes of early coral development under ambient pH (pH 7.8), and two low pH conditions (pH 7.6 and 7.3).

## Introduction

As anthropogenic carbon dioxide (CO_2_) levels continue to rise and contribute to global climate change (Bindoff et al., 2019), approximately one-third of atmospheric CO_2_ is also being absorbed by the surface ocean (Sabine et al., 2004). Increasing concentrations of CO_2_ in the surface ocean leads to ocean acidification (OA) through a shift in the chemical equilibrium of carbon in seawater (Sabine et al., 2004) that reduces ocean pH and the availability of carbonate ions (Caldeira and Wickett, 2003) for calcium carbonate formation. The absorption of atmospheric CO_2_ by the surface ocean has already driven a 0.1 unit decrease in oceanic pH, which is projected to decrease by an additional 0.036–0.042 or 0.287–0.29 units by the year 2100 under best (RCP 2.6) and worst-case (RCP 8.5) scenarios, respectively (Bindoff et al., 2019).

A growing body of studies has reported negative effects of reduced seawater pH on marine organisms. OA can negatively impact biomineralizing marine taxa that build calcium carbonate shells and skeletons (Doney et al., 2009; Kroeker et al., 2010). Coral reefs are one marine system threatened by OA as reef corals are ecosystem engineers, building skeletons that generate the 3-dimensional complexity and thus creating habitat for an entire ecosystem. Corals are also threatened by a variety of additional anthropogenic stressors such as marine heatwaves, pollution, sedimentation, and disease (Harvell et al., 2002; Hughes et al., 2017; Maynard et al., 2015). While adult organism calcification has been the primary response challenged by OA, less is known about the impact of OA on embryonic development, despite concerns that sensitivity to environmental stressors during early development can present a bottleneck for species persistence in a changing ocean (Byrne, 2012; Przeslawski et al., 2015). These early life stages are crucial for the resilience of species given their role in repopulating and replenishing existing populations (Adjeroud et al., 2017; Ritson-Williams and Arnold, 2009).

OA can affect multiple early developmental processes in corals and other marine invertebrates. Specifically, lower pH impacts sperm motility (Albright, 2011; Havenhand et al., 2008; Morita et al., 2010), fertilization success (Albright, 2011; Albright and Mason, 2013; Albright et al., 2010; Byrne, 2012; Han et al., 2021; Havenhand et al., 2008; Kurihara and Shirayama, 2004), larval settlement (Albright, 2011; Albright and Langdon, 2011; Albright et al., 2010), and post-settlement growth (Albright and Langdon, 2011; Albright et al., 2008; Albright et al., 2010; Scucchia et al., 2021a). In corals, decreased fertilization efficiency has been observed, which may be explained by decreased sperm flagellar motility and increased polyspermy under low pH (Albright and Mason, 2013; Albright et al., 2010). Low pH is also correlated with metabolic depression in many coral species (Moya et al., 2012; Moya et al., 2015; Nakamura et al., 2011; Putnam et al., 2013), which may explain observations of delayed development, smaller embryo/larval volume, and reduced post-settlement growth in corals developing in acidified conditions (Albright and Langdon, 2011; Nakamura et al., 2011; Rivest and Hofmann, 2014; Yuan et al., 2018). The increased energetic costs of maintaining homeostatic processes under disrupted acid-base balances and ionic gradients can additionally slow development and reduce growth of marine invertebrate larvae and recruits (Albright and Langdon, 2011; Byrne, 2012; Przeslawski et al., 2015; Timmins-Schiffman et al., 2013). For example, low pH is correlated to delayed cleavage (Foo et al., 2012), smaller larvae (Kelly et al., 2013), and depressed metabolism (O’Donnell et al., 2010) during the early development of sea urchins. Developmental delays (Barton et al., 2012; Timmins-Schiffman et al., 2013) and decreases in fertilization rate (Barros et al., 2013; Parker et al., 2009; Parker et al., 2010), hatching success (Barros et al., 2013; Gazeau et al., 2010; Van Colen et al., 2012), larval survival (Barros et al., 2013; Van Colen et al., 2012) and larval size, (Barros et al., 2013; Barton et al., 2012; Van Colen et al., 2012) have also been observed in bivalves under low pH. These compounding effects of low pH and co-occurring stressors can present a major life history bottleneck, leading to downstream effects on the resilience of marine invertebrate species (Byrne, 2012; Byrne and Przeslawski, 2013).

Most studies examining the effects of OA on coral development thus far have focused primarily on fertilization, settlement, and post-settlement growth, with less of a focus on embryonic development. In many species, embryogenesis represents a critical window in development during which time embryos are particularly sensitive to environmental fluctuations outside of the expected range (Byrne, 2012; Byrne et al., 2009; Dahlke et al., 2020; Ericson et al., 2010; Ericson et al., 2012; Foo et al., 2012; Przeslawski et al., 2015). Several studies have implicated the loss of maternal defenses during the maternal-to-zygotic transition (MZT) as playing a role in increased environmental sensitivity during this period (Dahlke et al., 2017; Dahlke et al., 2020; Ericson et al., 2012; Foo et al., 2012; Przeslawski et al., 2015). The MZT describes the developmental reprogramming of gene expression from maternal to zygotic regulation (Tadros and Lipshitz, 2009; Vastenhouw et al., 2019). During the MZT, the maternally-provided gene products that regulate early development and protect embryos from expected stressors are degraded as zygotic transcription activates and intensifies (Tadros and Lipshitz, 2009; Vastenhouw et al., 2019). Higher mortality in response to environmental perturbations has been observed during this transition in fish gastrula (Dahlke et al., 2017; Dahlke et al., 2020), sea urchin blastula/gastrula (Ericson et al., 2012; Foo et al., 2012), and several other marine invertebrates (Przeslawski et al., 2015).

Maternal mRNA provisioning and the timing of the MZT in reef-building corals, thus far, have only been investigated in the Hawaiian rice coral, *Montipora capitata* (Chille et al., 2021). Under ambient conditions, the maternal mRNA complement in *M. capitata* eggs primarily provide housekeeping functions, including cell cycle, biosynthesis, transcription, signaling, and protein processing (Chille et al., 2021; Van Etten et al., 2020). However, DNA repair mechanisms, which may protect early embryos from ultraviolet (UV) radiation and reactive oxygen species (ROS), are also over-represented in the *M. capitata* maternal mRNA complement of eggs (Chille et al., 2021). As of yet, the maternal mRNA complement in reef-building corals has not been assessed under stressful conditions. However, based on timing of the MZT in *M. capitata* (Chille et al., 2021), and general developmental gene expression studies in *Acropora millepora* (Strader et al., 2018), *Acropora digitifera* (Reyes-Bermudez et al., 2016), and *Nematostella vectensis* (Helm et al., 2013), coral embryos may be most sensitive to stress during early gastrulation (between 9 and 14 hours post-fertilization; hpf), when a large portion of maternal transcripts are degraded and zygotic transcription begins.

Previous transcriptomic examinations of pH-stress during coral embryonic development have been done on aposymbiotic species (Acropora; Strader et al., 2018) and on post-settlement stages after the completion of the MZT (Moya et al., 2012; Scucchia et al., 2021a). Here, we use *Montipora capitata*, a hermaphroditic broadcast spawner with vertical symbiont transmission that is endemic to the Hawaiian archipelago (Heyward, AJ, 1986; Padilla-Gamiño and Gates, 2012) to examine the embryonic sensitivity to OA in the context of the MZT. In this study, we examined the physiological and transcriptomic outcomes of early coral development in five life stages from fertilization until the “completion” of early coral development (i.e. fully-developed larvae) under pH 7.8 (ambient), and two low pH conditions (7.6 and 7.3). This time series allows us to determine sensitivity to and phenotypic consequences of OA during and after the MZT.

## Materials and Methods

### Specimen collection

*Montipora capitata* egg-sperm bundles were collected under the Department of Land & Natural Resources Special Activity Permit 2018-50 from the reef adjacent to the Hawai‘i Institute of Marine Biology, Coconut Island, Kāne‘ohe Bay, O‘ahu Hawai‘i (21°25’58.0”N 157°47’24.9”W) at ∼21:00 on June 13, 2018. After collection, egg-sperm bundles were allowed to break apart. Eggs were separated from the sperm and rinsed 3 times with 0.2 uM filtered seawater, then snap-frozen and stored at -80 °C in ∼300 uL aliquots.

### Experimental set-up

Fertilization and early development occurred in nine 1.6 L conicals flow-through tanks set to pH 7.8 (ambient), pH 7.6, and pH 7.3 pH (n=3 conical tanks per treatment, Fig.1A). At 22 hpf, the embryos were transferred to replicate 118 ml bins with 152µm plankton mesh bottoms floating in 74L glass tanks (n=3 bins per tank, 2 tanks per treatment; Fig. 1B) Treatments were designed to reflect future ocean acidification conditions in coastal embayments (Jury et al., 2013; Shaw et al., 2016) including daily fluctuations naturally occurring in Kāne‘ohe Bay, Hawai‘i (Drupp et al., 2011). All treatments were controlled by a pH-stat feedback system in one header tank per treatment. Header tank conditions were continuously monitored using a Neptune Apex Reef Aquarium set-up (Energy Bar 832; Controller Base Unit; PM1 pH/ORP Probe Module; Extended Life Temperature Probe; pH Probe), and 99.99% food-grade CO_2_ was injected into the header tanks with gas flow solenoids (Milwaukee MA955) using inline venturi injectors (Forfuture-go G1/2 Garden Irrigation Device Venturi Fertilizer Injector) in a pressure driven line connected to a circulating pump (Pondmaster Pond-mag Magnetic Drive Water Pump Model 5) in each header tank (Fig. 1A). Water flow was delivered at a maximum potential flow rate of 7.57 liters per hour by 1.9 LPH (0.5 gallons per hour) Pressure Compensating Drippers for a conical turnover rate of once every ∼50 minutes, and once every ∼23 hours for a 74 L tank. The average light intensity, 96±8.823 mean±sem umol m^-2^ s^-1^ (n=24), was measured using a handheld PAR sensor (Underwater Quantum Flux Apogee instruments - Model MQ-510; accuracy = ±4%).

**Figure 1.**
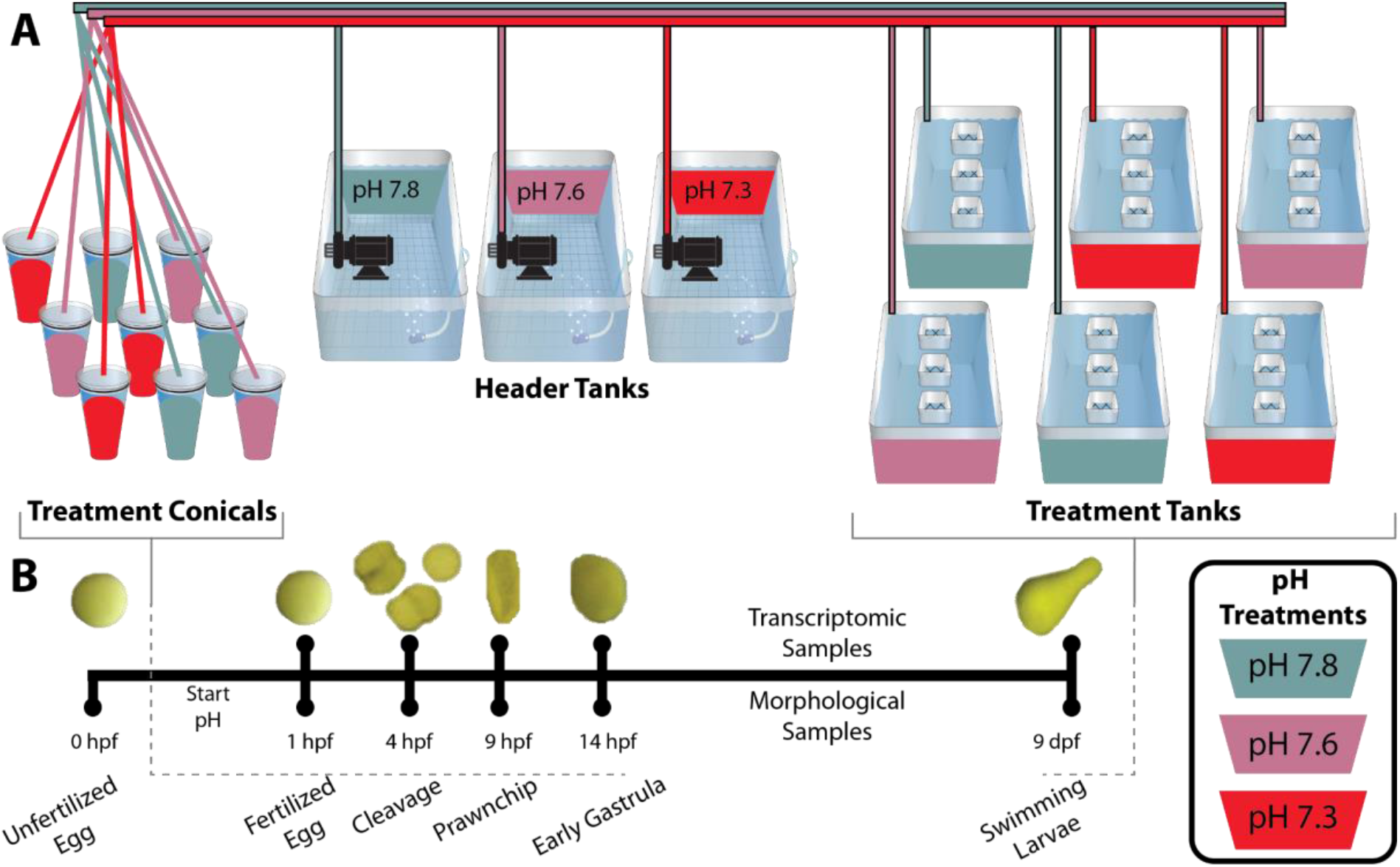
Summary of A) experimental design and B) sampling timeline, including example photographs of each life stage. Early life stages were exposed to treatment immediately after fertilization. Size analyses were performed on all life stages starting with Unfertilized Eggs (0 hpf). Gene expression analyses were performed on the samples following fertilization corresponding to fertilized eggs (1 hpf), cleavage (4 hpf), prawn chip (9 hpf), early gastrula (14 hpf) and swimming larvae (planula, 9 dpf).

Temperature, salinity, and pH were measured three times daily in each treatment tank using a handheld digital thermometer (Fisherbrand Traceable Platinum Ultra-Accurate Digital Thermometer, accuracy = ±0.05 °C, resolution = 0.001°) and a portable multiparameter meter (Thermo Scientific Orion Star A series A325; accuracy = ±0.2 mV, 0.5% of psu reading, resolution = 0.1 mV, 0.01 psu) with pH and conductivity probes (Mettler Toledo InLab Expert Pro pH probe #51343101; Orion DuraProbe 4-Electrode Conductivity Cell Model 013010MD), respectively (Table 1). Temperature and pH probes were calibrated using Tris (Dickson Laboratory Tris Batch 27, Bottles 70, 75, 167, 245, and 277) standard calibrations. Total alkalinity (µmol kg^−1^ seawater) was measured from water samples daily using an automated titrator (Mettler Toledo T50) and hydrochloric acid (Dickson Laboratory Titrant A3), and certified reference material (Dickson Laboratory CO_2_ CRM Batch 149) was used as a quality control standard. Carbonate chemistry parameters (pH, CO_2_, pCO_2_, HCO_3_, CO_3_, dissolved inorganic carbon, total alkalinity, and aragonite saturation) were calculated from total alkalinity measurements using SEACARB v3.0.11 (Gattuso et al., 2015) in RStudio (Table 1 and Table S1) using Kf from (Perez and Fraga, 1987), K1 and K2 from (Lueker et al., 2000) and Ks from (Dickson, 1990). Seawater pH was calculated on the total scale (Dickson et al., 2007); SOP6a), using the gas constant (8.31447215 J K^-1^ mol^-1^) and the Faraday constant 96485.339924 coulombs mol^-1^).

**Table 1.**
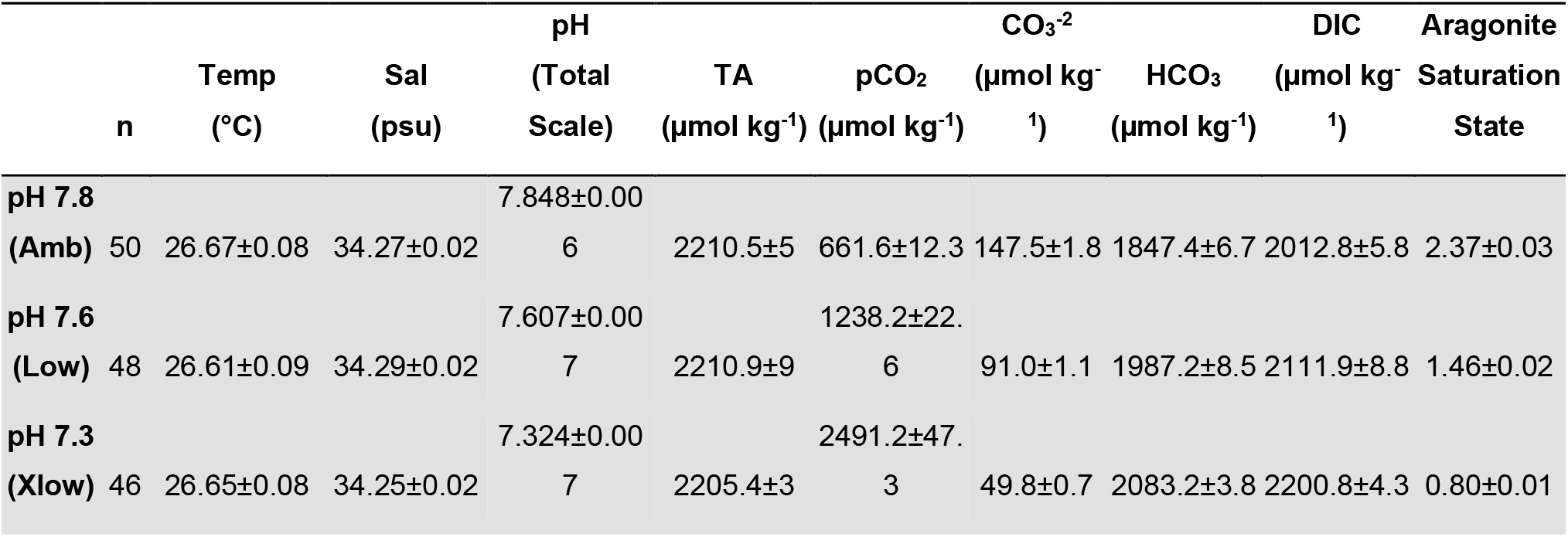
**Descriptive statistics for temperature, salinity, and carbonate chemistry across the 9 days of exposure (mean ± standard error of the mean).**

### Sample collection and preservation

Physiological and molecular samples were taken immediately upon breakup of egg-sperm bundles and at five timepoints following transfer to experimental conditions (Fig. 1). Sampling times represent visually-distinct developmental stages including fertilized embryo (1-hr post-fertilization), cleavage (4-hr post-fertilization), prawn chip (9-hr post-fertilization), early gastrulation (14-hr post-fertilization), and planula (9-d post-fertilization). Physiological samples (∼100-200 embryos or larvae per replicate) were preserved in 500 µL 25% zinc buffered formalin in 0.2µm filtered seawater, and stored at 4°C. Molecular samples (300 µL of embryos per replicate) were snap-frozen in liquid nitrogen and stored at -80°C.

### Embryo development and growth

Stereomicroscope images of fixed samples were analyzed for blastomere number at 4 hpf (cleavage stage), and for maximum length and perpendicular width of the unfertilized eggs, gastrula, and planula using ImageJ (Schneider et al., 2012). Cell division stage was assessed in cleavage-stage samples with 82 to 282 embryos/sample (n=3 replicate samples treatment^-1^).

Embryos were classified as “one cell,” “two cells,” “four cells,” or “greater than four cells.” The percentage of embryos at each stage of cell division was then calculated for each sample to examine potential variation in developmental timing between treatments. Volume of unfertilized eggs (n=50), fertilized embryos (n=225), early gastrula (n=200), and planulae (n=150) was calculated using the equation for an oblate ellipse: 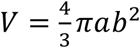 in which *V*is equal to volume, *a* is the half of maximum width and *b* is the half maximum length to examine differences in size between treatments through early development.

All statistical analyses for morphological data were performed in RStudio (v1.3.959; RStudio Team, 2020), using R version 4.0.2 (R Core Team, 2013). A beta regression model was used to analyze variation in the proportion of cells at each cleavage stage and treatment. Additionally, a one-way nested ANOVA analysis tested the effect of tank on embryo and planula size wherein tank ID was nested within treatment. After determining that tank effects were insignificant (P > 0.05), one-way ANOVA analysis was used to test for the effect of treatment on fertilized embryo, gastrula, and planula volume. Post hoc Tukey HSD tests were conducted when the effect of treatment was significant (P < 0.05). Data were visually examined for normal distribution and equal variance. The dependent variable was square-root transformed for the gastrulation and planula life stages in order to meet statistical assumptions prior to ANOVA analysis. Data points for these life stages were back-transformed for visualization in Figure 2.

**Figure 2.**
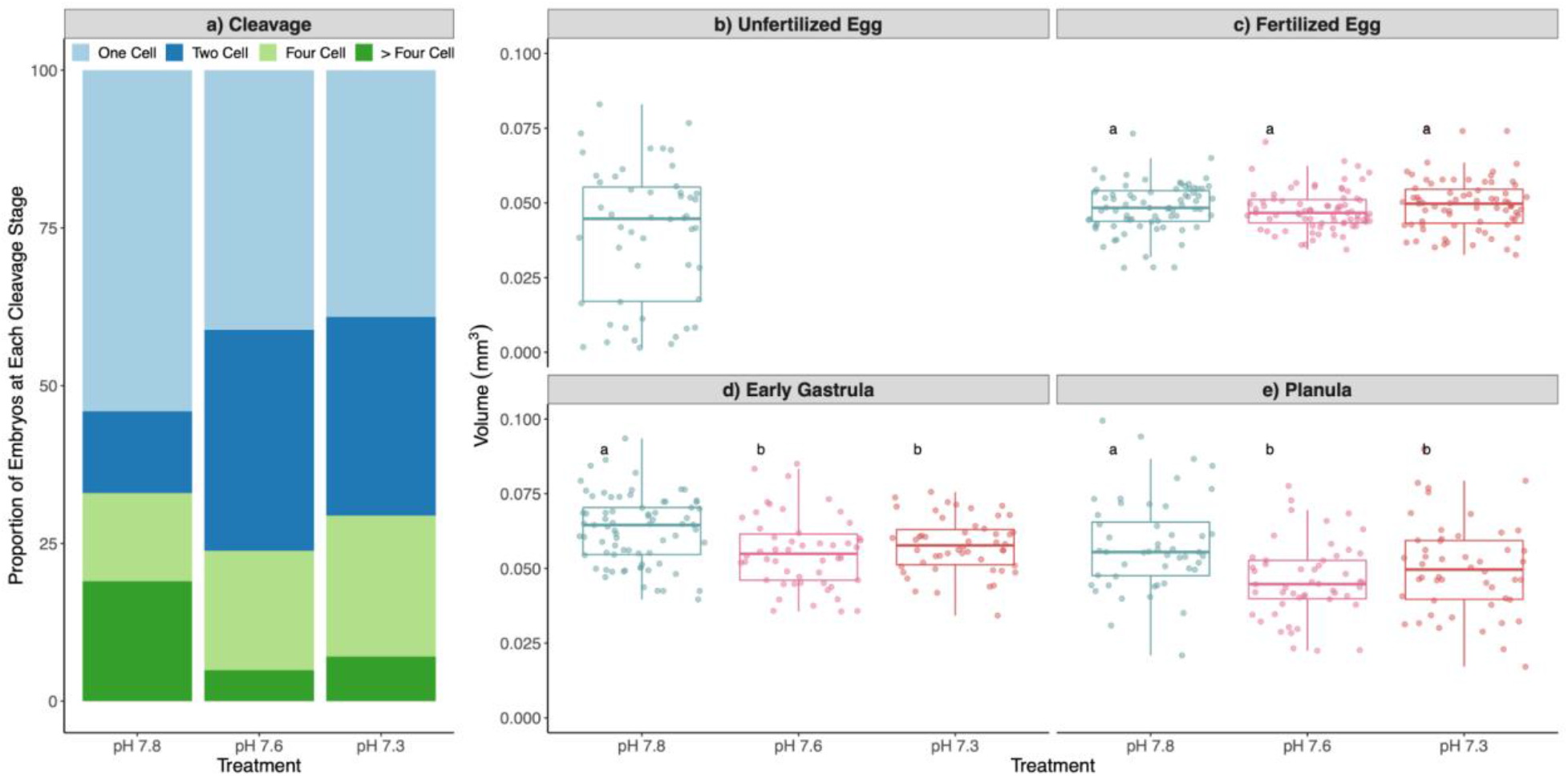
Developmental age and size. A) Proportion of embryos at each cleavage stage at 4 hpf and volume (mm^3^) of B) unfertilized eggs, C) fertilized eggs, D) early gastrula, and E) planula in each treatment. Prawn chip stages were not analyzed given the fold in the morphology which generates added assumptions for volume calculations (See Fig. 1 prawn chip picture).

### RNA extraction, sequencing, and processing

To extract RNA, molecular samples were digested at 55°C in 300 µl of DNA/RNA Shield (Zymo Research, Irvine, CA, USA) for two to three-and-a-half hours and centrifuged at 2200 RCF for one minute to separate the remaining solids. Total RNA was extracted from each supernatant with the Zymo Quick-DNA/RNA™ Miniprep Plus Kit (Zymo Research, Irvine, CA, USA) following the manufacturer’s protocol for tissue samples. RNA was quantified (ng/μL) with a ThermoFisher Qubit Fluorometer and quality was assessed using the Eukaryote RNA analysis on the Agilent TapeStation 4200 System. Total RNA samples were then sent to Genewiz (South Plainfield, New Jersey, USA) for library preparation and sequencing. cDNA libraries were constructed following the TruSeq Stranded mRNA Sample Preparation protocol (Illumina, San Diego, CA, USA) and sequenced on a HiSeq instrument targeting 15 million reads per sample. Quality trimming, alignment, and assembly were conducted as described in (Chille et al., 2021). In short, sequences were filtered for quality by applying a five base pair sliding window to remove reads with an average quality score less than 20, and retaining sequences with quality scores greater than or equal to 20 in at least 90% of bases and sequence length greater than or equal to 100 bases. Trimmed reads were then aligned to an *M. capitata* reference assembly (Shumaker et al., 2019) using HISAT2 in the stranded paired-end mode (Kim et al., 2019), and assembled with StringTie in the stranded setting (v2.1; Pertea et al., 2015). Finally, the StringTie *prepDE* python script (Pertea et al., 2015) was used to generate a gene count matrix from the resulting GFFs (Pertea et al., 2015).

### Host gene expression analyses

Experiment-wide patterns in gene expression were assessed with a principal components analysis (PCA). First, the full gene count matrix was pre-filtered improve to remove low-coverage counts with the Genefilter (v1.70.0; Gentleman et al., 2020) function *pOverA* to exclude genes where fewer than 2 out of 41 (∼0.0488) samples had counts under 10. A value of p=0.0488 was chosen under the assumption that due to the MZT, a gene count of zero likely represents a true zero because some genes in *M. capitata* embryos may not be activated at until the onset of gastrulation or later (Chille et al., 2021). Then, counts were transformed with DESeq2’s (v1.28.1) variance stabilizing transformation (vst) function (Love et al., 2014a) after confirming that all size factors were less than 4. Finally, the vst-transformed counts were used for PCA to visualize expression results for all filtered genes. The DESeq2 *plotPCA* function was used to generate a PCA of all samples based on sample-to-sample distance. All gene expression analyses were performed in RStudio (v1.3.959; RStudio Team, 2020), using R version 4.0.2 (R Core Team, 2013).

PCA and differential gene expression analysis with DESeq2 were used to assess within-stage differences in expression due to treatment. As described above, genes with low overall expression were filtered using the Genefilter *pOverA* filter function (Gentleman et al., 2020). After testing a range of values from p=0.33 to p=0.9, a more conservative value of p=0.875 was chosen to prevent bias due to high duplication levels in the first four life stages where zeros likely represent true zeros compared to the planula samples which had lower duplication. As such, we retained genes with greater than 10 counts in at least 7 out of 8 samples (pOverA 0.875, 10) for further analysis. The resulting genes were then vst-transformed using the DESeq2 *vst-transformation* function (Love et al., 2014b). PCA was conducted with the DESeq2 *plotPCA* function (Love et al., 2014b), using vst-transformed counts to calculate sample-to-sample distance. Differential expression analysis between pH treatments was then performed for each life stage using the filtered counts (pOverA 0.875, 10). Pairwise differences in gene expression between treatments were estimated using Wald model in DESeq2 (Love et al., 2014b). As DESeq2 results showed zero differentially-expressed genes (DEGs) in all treatment comparisons prior to the planula stage, only genes with padj<0.05 in the planula stage were used for further analysis. To visualize the DEGs that were shared and distinct for each treatment comparison within the planula stage, the results were summarised as a venn diagram (Fig. 4A) using the VennDiagram R package v1.6.20 (Chen and Boutros, 2011). Overall expression patterns among the planula DEG set were visualized as a heatmap using the ComplexHeatmap package v2.5.4 (Gu et al., 2016), with clustering based on differences in expression compared to the rowmean in the vst-transformed count matrix. Clusters in the heatmap are based on k-means clustering using the R package NbClust v3.0 (Malika et al., 2014) to calculate the optimal number of clusters using 30 indices.

### Host gene ontology enrichment analyses

Gene ontology (GO) enrichment analysis was performed on the planula DEG set to examine the biological processes and molecular functions primarily correlated with each DEG k-means cluster. Firstly, gene ontology annotation of the *M. capitata* genome was retrieved from (Chille et al., 2021). To supplement the GO annotation for functional enrichment analysis, Kegg annotation was completed using KofamScan (v1.3.0; Aramaki et al., 2020), to retrieve Kegg terms in the eukaryote profiles database (v. April 23, 2021). GO enrichment analysis was performed on the DEGs with log2FoldChange>|1| in each k-means cluster using the R package Goseq (v1.40.0; Young et al., 2010). Subsequently, slim categories for the enriched GO terms were obtained using the *goSlim* function in the R package, GSEABase, using the generic GOslim obo database as reference (http://current.geneontology.org/ontology/subsets/goslim_generic.obo; v1.2; Ashburner et al., 2000). Finally, enriched biological process and molecular function terms with numInCat > 5 and their associated slim categories were visualized as a heatmap using the ggplot2 *geom_tile* function (Wickham, 2011).

### Symbiont transcriptome assembly and gene expression analysis

To assess correlation between pH Treatment and gene expression of Symbiodinaceae within the planula host, the symbiont transcriptomes were extracted from a *de novo*-assembled holobiont sequence library and analysed for differential expression between treatments. The holobiont transcriptome was assembled *de novo* using Trinity v2.12.0 (Grabherr et al., 2011) with the stranded ‘RF’ option. The 9 planula libraries were used to construct the holobiont transcriptome. The three adult *Montipora capitata* libraries from (Chille et al., 2021) were additionally used for *de novo* assembly in order to obtain a symbiont-rich library. Following assembly, completeness was assessed using BUSCO v4.0.6 (Simão et al., 2015) to detect the presence of single-copy orthologs universal to alveolata (for the symbionts) and metazoans (for the host). The host and symbiont transcriptomes were then identified from the holobiont assembly using *psytrans* (https://github.com/sylvainforet/psytrans) using *Cladocopium* C1 (Liew et al., 2016; Voolstra et al., 2015) and *Montipora capitata* (Shumaker et al., 2019) predicted proteins as references.

Following host-symbiont separation, the symbiont reads were used as a reference file for the Trinity *align_assemble* perl script (Grabherr et al., 2011) using RSEM v1.3.3 (Li and Dewey, 2011) for abundance estimation and Bowtie v-1.2.1.1 as the alignment method (Langmead, 2010). Finally, the symbiont gene counts were obtained using the Trinity *abundance_estimates_to_matrix* perl script (Grabherr et al., 2011). Differential gene expression of the planula endosymbionts was carried out using the methods described for the host early life stages (see *Differential gene expression analysis*). Finally, the identities of the symbiont DEGs were determined through a local alignment search of the corresponding sequences against the NCBI non-redundant database using Blastx v2.11.0 (Altschul et al., 1997) with default parameters.

## Results

### Embryo development and growth

Immediately after release, prior to treatment exposure, eggs were 0.040±0.023 mm^3^ (Fig. 2A). Following treatment exposure, the proportion of embryos at each cleavage stage at 4 hpf and embryo size in the fertilized egg (1 hpf), early gastrula (14 hpf), and planula (9 days post-fertilization; dpf) samples was assessed to characterise morphological effects of developing in ambient (7.8) low (7.6) and extreme low pH (7.3) conditions. At the cleavage stage, across all treatments combined undivided cells were on average more frequent (44.77 ± 21.49%; mean ± sd) than those at the two-cell (26.51 ± 12.63%), four-cell (18.37 ± 11.23%), or greater-than-four cell stage (10.34 ± 13.40%). Beta regression analysis of the proportion of cells at each cleavage stage and treatment (Fig. 2B) shows that while there were significant differences (<0.0001) in proportions of two-, four-, and greater-than-four cell stages overall, the interaction between cleavage stage and treatment were only marginally significant (p=0.0765). Additionally, there were no significant differences between pH treatments in embryo size in the fertilized eggs (p=0.256; Fig. 2C). However, at both the early gastrula (p=0.000292; Fig. 2D) and planula stages (p=0.000109; Fig. 2E) embryos developing under ambient pH conditions were significantly larger than those developing under pH 7.6 and 7.3 conditions.

### RNA sequencing, quality control, and mapping

TruSeq Illumina Stranded Paired-End (PE) sequencing resulted in 935,000,000 PE (19,392,156 mean reads/sample), with 694,800,000 PE reads (14,241,176 mean reads/sample) remaining after quality filtering and adapter removal. On average, 79.28 ± 1.95% (mean ± sem reads per sample) aligned to the reference *M. capitata* genome (Shumaker et al., 2019). Assembly quality assessment with GFFcompare showed that 63,227 genes queried from the mapped GTF files were matched to the 63,227 genes in the reference genome (Shumaker et al., 2019), with 40,883 matching intron chains, 63,218 matching loci, and zero missing elements.

### Host gene expression patterns

After filtering to remove genes with fewer than 10 counts in ∼4.88% of samples (i.e. the proportion representing each treatment in a given life stage), 30,271 genes remained to characterize global gene expression patterns. A principle components analysis (Fig. 3A) shows that sample-to-sample variation is primarily related to differences in life stage, with PC1 (83%) and PC2 (12%) explaining the majority of variation between life stages.

**Figure 3.**
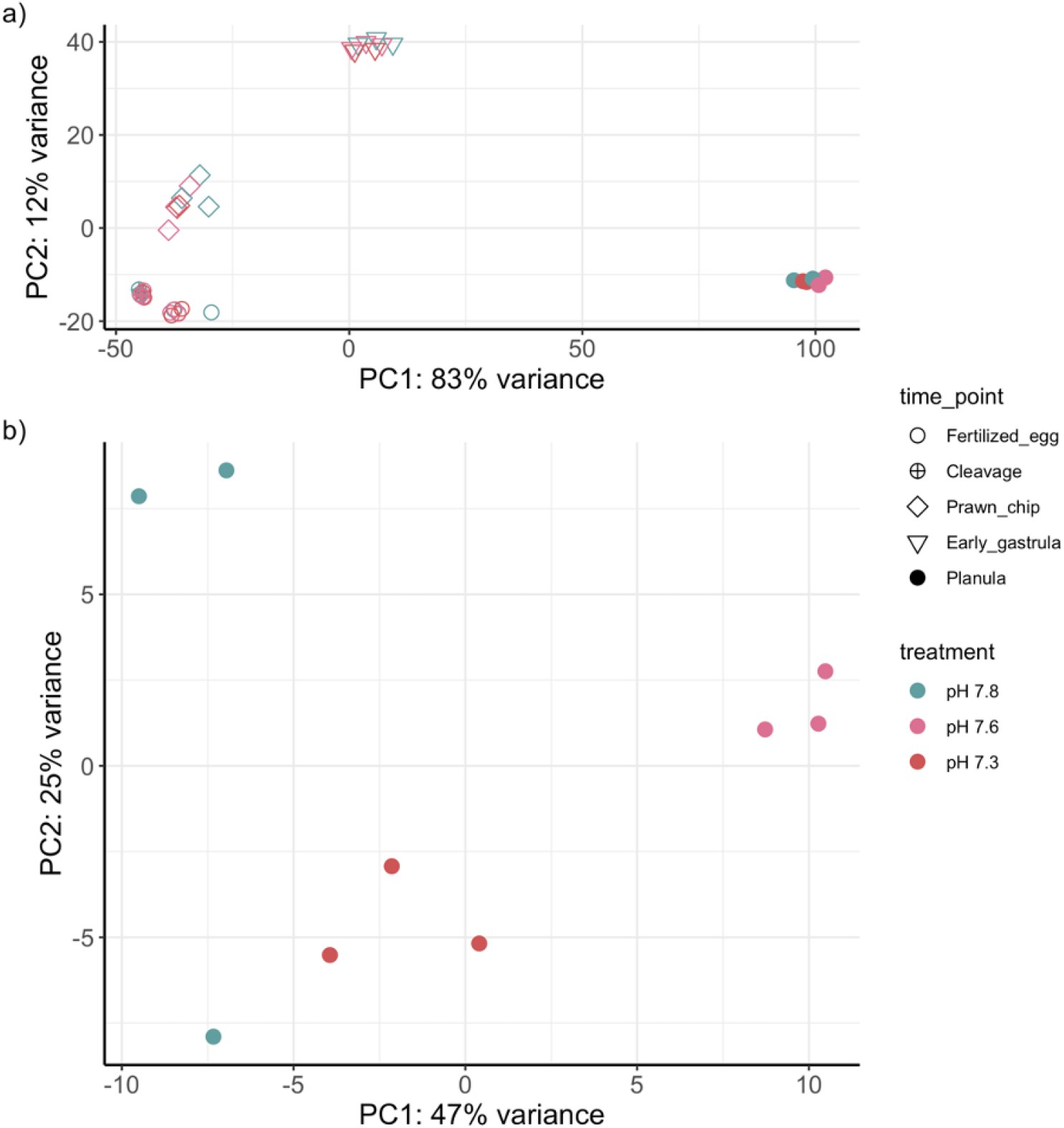
Principal components analyses of sample-to-sample distances based on vst-transformed gene expression. A) principal components analysis of all life stages based on sample-to-sample distance computed from all genes passing a low counts filter, wherein a gene must have a count of 10 or greater in at least 2 out of the 41 samples (pOverA ∼0.05, 10). B) shows principal components analysis of the sample-to-sample distance of planula samples, only (pOverA ∼0.875, 10).

Principal components analyses computed from variation in expression of genes with greater than ten counts in seven of eight fertilized egg (10,751 genes; Fig. S1A), cleavage (12,837 genes; Fig. S1B), prawn chip (16,392 genes; Fig. S1C), early gastrula (17,669 genes; Fig. S1D), and planula (23,225 genes) samples show no differentiation between treatments until the planula stage (Fig. 3B). At the planula stage, treatment groups separate strongly along PC1 and PC2, which explain 47% and 25% of the variation between samples, respectively. Notably, ambient pH (7.8) and extreme low pH (7.3) samples cluster more similarly on PC1 than ambient and low pH (7.6) samples. However, with the exception of one ambient sample, ambient and low pH samples cluster closer than ambient and extreme low pH samples on PC2 (Fig. 3B).

Pairwise treatment comparisons within each life stage using DESeq2 (Table S2) showed zero differentially expressed genes between treatments until the planula stage. Differentiation of samples from planula raised in different treatments are supported by 5,406 DEGs between planula samples raised under the two low pH conditions relative to the ambient pH conditions. Mirroring the clustering of treatments along PC1 in Fig. 3B, a venn diagram (Fig. 4A) of shared and unique DEGs in each treatment comparison shows 2,202 DEGs between planula samples at the low and ambient pH treatments, 1,078 of which are also differentially expressed between the extreme low and low treatments. Only 66 DEGs exist between planula samples at the extreme low and ambient pH treatments. Planula DEGs were grouped into two k-means clusters based on the consensus of 11 of 30 clustering indices (Fig. 4B), with genes in Cluster 1 upregulated in the low pH treatment compared to the ambient and extreme low pH treatments, and genes in Cluster 2 exhibiting the opposite pattern (Fig. 4B).

**Figure 4.**
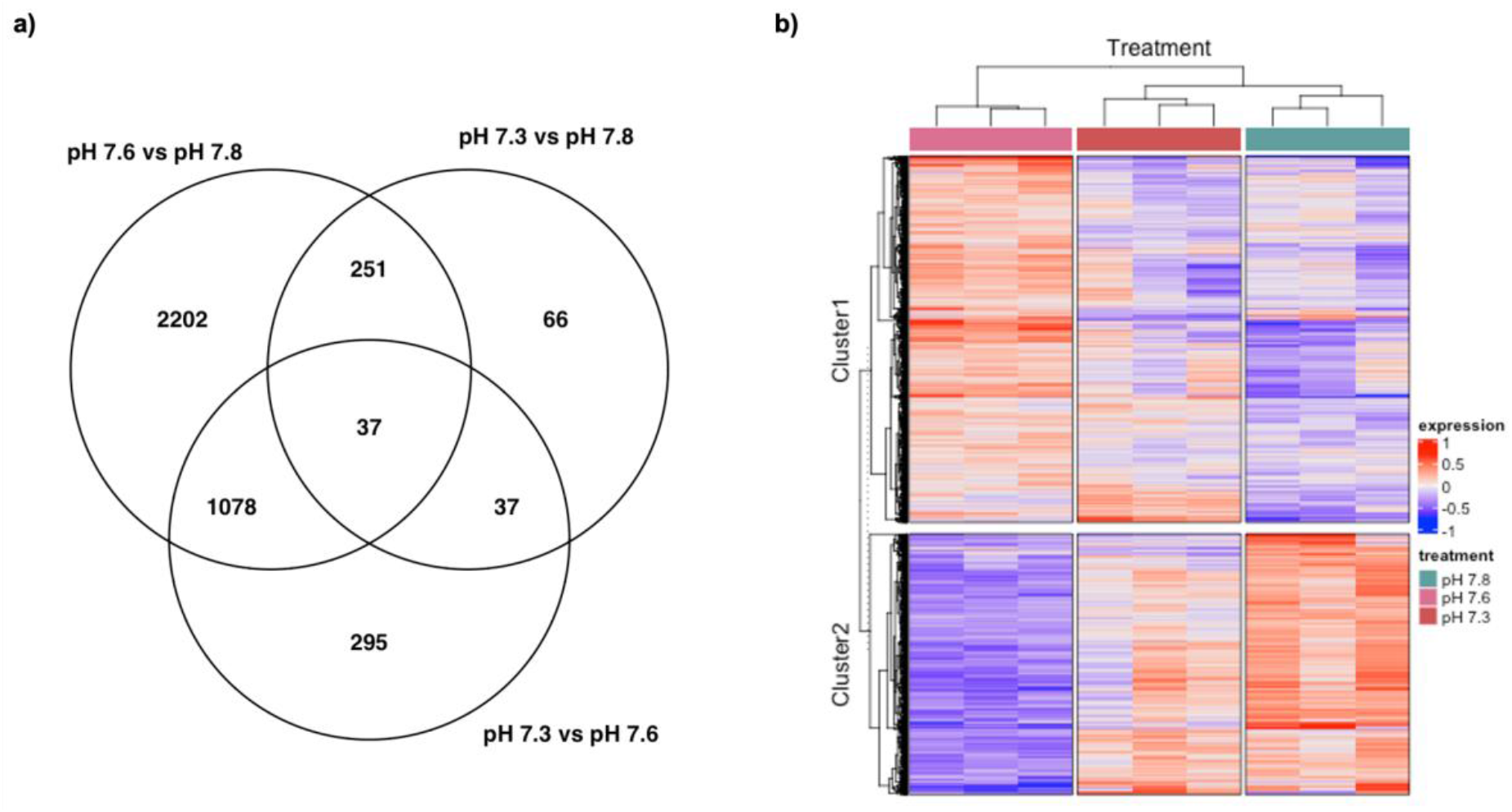
A) Venn diagram and B) heatmap of all genes that are differentially-expressed between treatments at the planula stage. Clusters in the heatmap are based on k-means clustering using the R program NBclust to calculate the optimal number of clusters using 30 indices. Heatmap colors for each gene are based on difference in expression compared to average VST across all samples, where red cells are more highly expressed compared to the rowmean and blue cells more lowly expressed compared to the rowmean.

### Host gene ontology enrichment

After filtering the DEG set to retain genes with log2FoldChange>|1|, 344 DEGs remained in Cluster 1 (upregulated under low pH), and 256 remained in Cluster 2 (downregulated under low pH) for GO enrichment analysis. GO enrichment analysis revealed 39 biological processes and 31 molecular functions over-represented in Cluster 1. Biological processes overrepresented in Cluster 1 (Fig. 5A, Table S5) are primarily related to 7 slim terms, including biosynthetic process, cellular component assembly, cellular nitrogen compound metabolic process, immune system process, lipid metabolic process, response to stress, and transport. Likewise, molecular functions overrepresented in Cluster 1 (Fig. 5B, Table S5) can be summarised by 8 slim terms, including ATPase activity, cytoskeletal protein binding, DNA binding, ion binding, kinase active, oxidoreductase activity, structural molecule activity, and transferase activity. Ion binding, in particular, includes several offspring terms under the GO category calcium ion binding. In Cluster 2, GO enrichment revealed 5 overrepresented biological processes (Fig. 5A, Table S5) represented by the slim terms biosynthetic process, immune system process, and transport, as well as 10 overrepresented molecular functions (Fig. 5B, Table S5) represented by the slim terms, ATPase activity, nucleotidyltransferase activity, peptidase activity and transmembrane transporter activity. The GO terms enriched in Cluster 1 and Cluster 2 are supported by the overrepresentation of related Kegg terms in both clusters (Table S6), such as peroxidase [K19511], Amt family ammonium transporter [K03320], cytosolic phospholipase A2 [K16342], and calreticulin [K08057] in Cluster 1 and E3 ubiquitin-protein ligase [K22754] and [histone H4]-lysine20 N-methyltransferase SETD8 [K11428] in Cluster 2.

**Figure 5.**
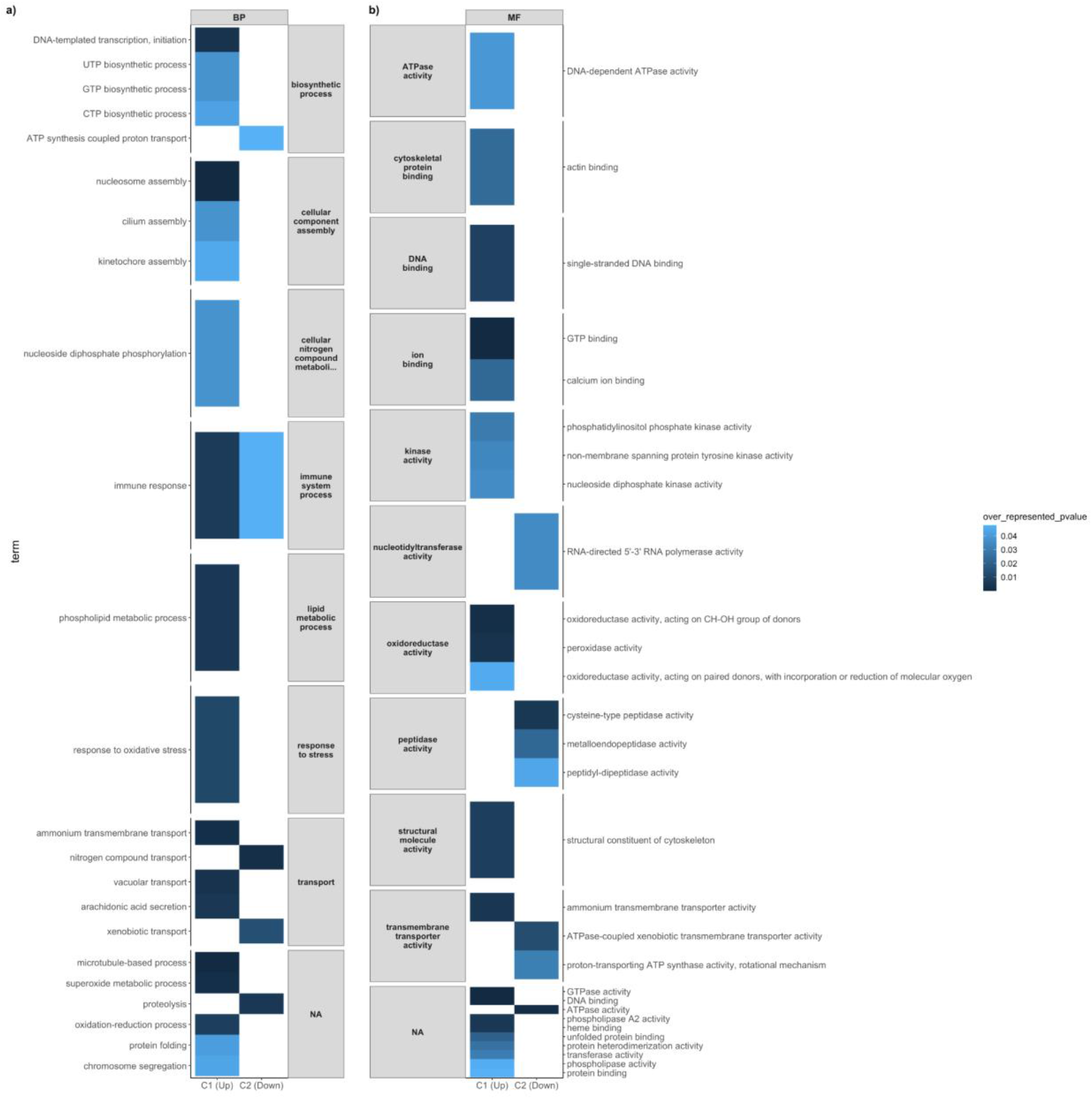
a) Biological process and b) Molecular Function terms that are enriched in planula developing in the low treatment compared to the extreme low and ambient treatments. Up = Terms associated with DEGs (p-adjusted<0.05, log2FoldChange>|1|, numInCat>5) from Cluster 1 (C1), Down = terms associated with DEGs (p-adjusted<0.05, log2FoldChange>|1|, numInCat>5) from Cluster 2 (C2)

### Symbiont transcriptome assembly and gene expression

The *M. capitata* planula and adult holobiont *de novo* assembly was constructed from 698,861,565 bases, yielding 991,330 transcripts and 656,603 genes. The N50 of the holobiont assembly was 1,102, with an average contig length of 704.97 bases and a GC content of 43.7%. A BUSCO search of the holobiont assembly against the metazoan and alveolata databases for the host and symbiont showed high completeness (90.1% alveolata and 97.5% metazoan). In total, 83.37% (826,513) of transcripts from the holobiont assembly were assigned to the host and 16.63% (164,817) of transcripts were assigned to the symbiont. Finally, using the separated assemblies as reference transcriptomes for Trinity, 4.77±0.43% (mean±se) of planula sequences aligned to the symbiont, yielding 133,811 symbiont genes for differential expression analysis.

After filtering to remove reads with fewer than 10 counts in 7 out of 8 samples, 1,365 genes remained for principal components and differential expression analyses. A principle components analysis (Fig. S2A) of the symbiont samples shows large variation in gene expression between the three low pH (7.6) samples along the PC1 (28%) and PC2 (14%) axes, but the samples from the ambient pH (7.8) and extreme low pH (7.3) each cluster by treatment. Differential gene expression analysis with DESeq2 (Table S3) revealed 12 differentially-expressed genes (DEGs) between the ambient and low treatments, 17 DEGs between the ambient and extreme low treatments, and 0 DEGs between the low and extreme low treatments. The ten symbiont genes upregulated in the low treatment compared to the ambient aligned to the histone H3 protein as well as actin and tubulin proteins, while the two downregulated genes aligned to the pre-mRNA-splicing factor ATP-dependent RNA helicase and a cell wall-associated hydrolase (Table S4). However, the seventeen symbiont genes upregulated in extreme low treatment (Table S4) compared to the ambient were primarily associated genes regulating rDNA expression, a mucin-like protein, cytochrome P450, COPB protein, as well as several hypothetical proteins.

## Discussion

Limited research has been conducted so far on the effects of OA on coral embryonic development, with a primary focus on physiological processes (Chua et al., 2013a; Chua et al., 2013b), while little is known about the changes occurring at the transcriptomic level. Here we show that while acidic seawater altered the morphological development of *M. capitata* embryos, it did not drive any substantial change in gene expression until after the MZT. Early gastrula and planula developing at pH 7.6 (low) and pH 7.3 (extreme low) were significantly smaller than those developing at pH 7.8 (ambient), suggesting an energetic cost to developing under low pH. This is also supported by functional enrichment of the genes associated with cell division, regulation of transcription, lipid metabolism, and oxidative stress that are significantly upregulated in swimming larvae (9 dpf) exposed to pH 7.6. We additionally show that transcriptomically, larvae developing at pH 7.8 and pH 7.3 were more similar than those developing at pH 7.6 and hypothesize that this may be due to high CO_2_ stimulation of symbiont photosynthesis at pH 7.3, which may alleviate the effects of OA on the host fitness.

### The MZT under ocean acidification

While the morphological effects of low pH were apparent by the early gastrula stage (22 hpf; Fig. 2D-E), pH did not drive any substantial changes in gene expression until the larval stage, i.e., after the zygotic transcriptome has assumed autonomy (Fig. S1; Chille et al., 2021). These results suggest that the maternal-to-zygotic transition in *M. capitata* is robust to low pH. While our study does not include survivorship data, this finding is supported by other studies on OA stress during the embryonic development of fish (Dahlke et al., 2017; Dahlke et al., 2020) and sea urchins (Ericson et al., 2012; Foo et al., 2012) that show that while early development is robust to ocean acidification, embryos are more vulnerable after the MZT. These findings support the role of maternal gene products (i.e., mRNA, proteins, transcription factors) in buffering early embryos from environmental stress (Hamdoun and Epel, 2007). In *M. capitata*, early development is likely to be influenced by maternal transcripts until 22-36 hpf, when the zygotic transcriptome exhibits the capacity for responding to stimuli (Chille et al., 2021). Seeing as a transcriptomic response to decreased pH was only apparent at the planula stage (Fig. 3b & Fig. 4), this suggests that the transcripts necessary to buffer embryonic development from OA are already expressed under ambient conditions or that embryos possess other molecular defenses, such as maternally-provisioned proteins (Hamdoun and Epel, 2007) that buffer the effects of low pH. However, maintaining homeostasis under low pH comes with an energetic cost, as indicated by the smaller sizes of embryos developing low pH. As such, while the embryonic development of *M. capitata* may be robust to ocean acidification, the energetic cost of developing under low pH may have latent effects on larval competence and recruitment (Albright, 2011).

### Outcomes of developing in low pH

In our study, low pH (pH 7.6) elicited a robust transcriptomic response in planula that implicates several processes affecting energy metabolism and larval size (Fig. 2E), including biomineralization, cellular acidosis, and cellular stress response. We detail these responses below.

Several studies have found that actively calcifying larvae may be more susceptible to changes in seawater carbonate chemistry compared to non-calcifying embryos and larvae (Byrne, 2012; Przeslawski et al., 2015) by making biomineralization more chemically and energetically difficult (Ries, 2011). In the brooding coral *Stylophora pistillata*, calcium carbonate deposition is initiated during the pre-settled larval phase (Akiva et al., 2018). While mineralization was not specifically examined in our study, as nine-day-old larvae, it is likely that the planula here were at the calcifying stage. In fact, gene ontology enrichment analysis shows that genes associated with calcium ion binding and calcification, such as calreticulin, are upregulated in larvae exposed to pH 7.6 (Table S5). Genes involved in calcium ion binding like calreticulin provide structural and cell adhesion functions that aid in the biomineralization process (Mass et al., 2016) and may be upregulated to compensate for the chemical shift in carbon equilibrium in the seawater (Comeau et al., 2018; DeCarlo et al., 2018; Zhao et al., 2020). However, upregulation of these genes in response to OA may increase energetic demand, as observed in some adult corals (Davies et al., 2016; Vidal-Dupiol et al., 2013; Scucchia et al., 2021b). This increased demand could potentiate the rapid depletion of larval lipid stores if the energy derived from their endosymbionts is insufficient to keep up with demand (Alamaru et al., 2009; Ben-David-Zaslow and Benayahu, 2000; Harii et al., 2010), ultimately decreasing larval size.

Acid-base parameters such as pH and pCO_2_ are generally considered to lead to depressed metabolism through cellular acidosis, leading to alterations in cell size (Kaniewska et al., 2012; Reipschläger and Pörtner, 1996). Here, the enrichment of terms related to ATP synthesis in the genes downregulated at low pH (pH 7.6; Table S5), and lipid metabolism in the genes upregulated at low pH (pH 7.6; Table S5) suggests alteration of larval metabolism at reduced pH. Metabolism is highly interconnected with both growth and cell cycle progression (Björklund, 2019). Additionally, mitochondria play a crucial role in setting growth rates through energy generation and lipid biosynthesis (Miettinen and Björklund, 2017). In this context, alteration in metabolism at low pH might cause changes in cell size. Here, the enrichment of terms related to cell growth and cytoskeleton in the genes upregulated at low pH 7.6 (pH 7.6; Table S5) support changes in cell cycle and cell size with acidification. Cell growth-related genes such as JNK cascade and GTPase activity have been shown to change with acidification (Liew et al., 2018), and acidic cellular pH was demonstrated to alter processes such as cytoskeletal integrity (Tominaga and Barber, 1998). Such alterations in cytoskeletal integrity could explain the observed smaller size of planula that developed under reduced pH conditions.

Chronic exposure to low pH (pH 7.6) during early development also appears to elicit the onset of cellular stress and antioxidant responses. For example, GO terms enriched in planula developing at pH 7.6, including oxidative stress, immune response, superoxide metabolic process, oxidation-reduction process, and microtubule-based process (Table S5) are linked to protein domains that protect the cell from oxidative stress through oxidoreductase activity (e.g., superoxide dismutase-like, peroxidasin-like, and FAD-linked oxidase; DeSalvo et al., 2008; DeSalvo et al., 2010; Fernandes et al., 2007; Kaniewska et al., 2012; Lesser and Farrell, 2004; Martínez, 2007; Sampayo et al., 2003; Zhao et al., 2016) and the removal of damaged cells (e.g., Toll/interleukin-1 receptor, TNF receptor-associated factor 4; Fuchs and Steller, 2015; Linkermann and Green, 2014; Poquita-Du et al., 2019; Todgham and Hofmann, 2009; van de Water et al., 2015).

Upregulation of cellular stress and antioxidant responses is consistent with stress response mechanisms documented in adult and larval corals upon exposure to multiple stressors, including low pH (Kaniewska et al., 2012), ultraviolet radiation (Lesser and Farrell, 2004), and thermal (DeSalvo et al., 2008; DeSalvo et al., 2010; Polato et al., 2013; Rosic et al., 2014), and nutrient stress (Rosic et al., 2014). Here, cellular stress and antioxidant responses may have been induced from macromolecular damage due to ROS released from the rapid peroxidation of planula lipid stores for energy use (Polato et al., 2013) or from increased ROS production by the algal endosymbiont. Notably, however, the onset of cellular stress response may also come at an energetic cost to the planula (Sokolova et al., 2012) and lead to smaller larvae by depleting maternal lipid stores.

### Symbiont stimulation under high CO_2_

Predicted future increase of CO_2_ in seawater has been shown to stimulate the photosynthetic activity of the algal symbiont (Krief et al., 2010; Reynaud et al., 2003). Such stimulation can enhance host fitness under acidification (Guillermic et al., 2021). In this study, the enrichment of terms related to coral oxidative stress under pH 7.6 (low; Table S5) suggests increased photosynthetic activity in the endosymbiotic algae due to the higher availability of CO_2_ in seawater, as observed in the endosymbionts of *Stylophora pistillata* primary polyps and adults (Scucchia et al., 2021a; Scucchia et al., 2021b). It has been hypothesized that at higher concentrations of CO_2_ the stimulation of symbiont photosynthesis could prevent acidosis of the host cells, thus helping with preserving cellular acid-base homeostasis (Gibbin et al., 2014), and ultimately alleviating the effects of OA on the host fitness, as observed here in planula developing at pH 7.3. The increase in translocated photosynthates provided by the endosymbiont under this scenario could additionally supplement the larvae’s diet (Harii et al., 2010), thereby decreasing consumption of endogenous lipids and the associated influx of ROS. The differential expression of housekeeping functions, including tubulin, rRNA promoter binding proteins, and rDNA transcriptional regulator 15 in our symbiont data (Table S3) suggest to us that there is not enough endosymbiont expression coverage in the dataset to have the power to test this hypothesis. It is possible, however, that due to the potential for post-transcriptional regulation of gene expression in the Symbiodiniaceae through trans-splicing of spliced leader sequences (Zhang et al., 2007; Lin et al., 2010; Lin, 2011), small RNAs and microRNAs, (Baumgarten et al., 2013; Lin et al., 2015), and/or RNA-editing (Liew et al., 2017), differential expression may not necessarily be observed.

### Conclusions

In this study, we examined developmental sensitivity to and phenotypic consequences of OA during and after the MZT in the rice coral, *Montipora capitata*. While on a transcriptomic level, no differential gene expression was observed in the host until after the MZT, phenotypically, early gastrula (14 hpf) and swimming larvae developing in the pH 7.6 and pH 7.3 treatments were significantly smaller than those in the control pH 7.8. This supports that the development of *M. capitata* embryos is resilient to low pH conditions, even those substantially lower than what currently occurs in Kāneʻohe Bay. While the MZT appears to be robust to OA, the energetic strain on larvae from maintaining homeostasis under low pH could create negative carryover effects for the energetically costly process of metamorphosis (Edmunds et al., 2013) and may make recruits more susceptible to competition and predation (Anlauf et al., 2011; Pechenik et al., 2002; Thiyagarajan et al., 2007). Lowered fitness in these early life stages could potentially create an ontogenetic bottleneck that impacts the repopulation and replenishment of coral reefs and contribute to their continued decline (Hughes and Tanner, 2000). Of note, however, the reductions in body size detected so far in coral larvae represent the effects of future OA conditions on modern populations without consideration of potential moderating effects of rapid adaptation or acclimatization, which is pivotal to accurately forecast the impacts of climate change on marine organisms (Kelly et al., 2013; Putnam, 2021).

## Acknowledgements

We would like to extend our gratitude to the faculty and staff of the Hawaiʻi Institute of Marine Biology and the University of Rhode Island Computing for accommodating us and for their technical support.

## Data Availability

All raw data can be accessed on the NCBI SRA repository under the Bioproject accession IDs, [PRJNA731596](https://www.ncbi.nlm.nih.gov/bioproject/731596) (adult) and PRJNA616341 (all other timepoints). Additionally, all scripts used for bioinformatic and statistical analyses are available on the GitHub repository, Mcapitata_OA_Developmental_Gene_Expression_Timeseries.

## List of Abbreviations

CO_2_: carbon dioxide
OA: ocean acidification
MZT: maternal-to-zygotic transition
UV: ultraviolet
ROS: reactive oxygen species
hpf: hours post-fertilization
LPH: liters per hour
dpf: days post-fertilization
vst: variance stabilizing transformation
PCA: principal components analysis
GO: gene ontology
DEG: differential-expressed gene
PE: paired-end

