## Supplementary_Tables_and_Figures for "Energetics but not development is impacted in coral embryos exposed to ocean acidification"

---

### Supplementary Tables and Figures

**Table S1.** Discrete measurements for temperature, salinity, and OA chemistry corresponding to Table 1 (link)

**Figure S1.** Principal coordinates analysis of A) fertilized eggs, B) cleaving embryos, C) prawn chip, and D) early gastrula based on sample-to-sample distance computed from genes passing a low counts filter, wherein a gene must have a count of 10 or greater in at least 7 out of 8 samples (pOverA ~0.875, 10).

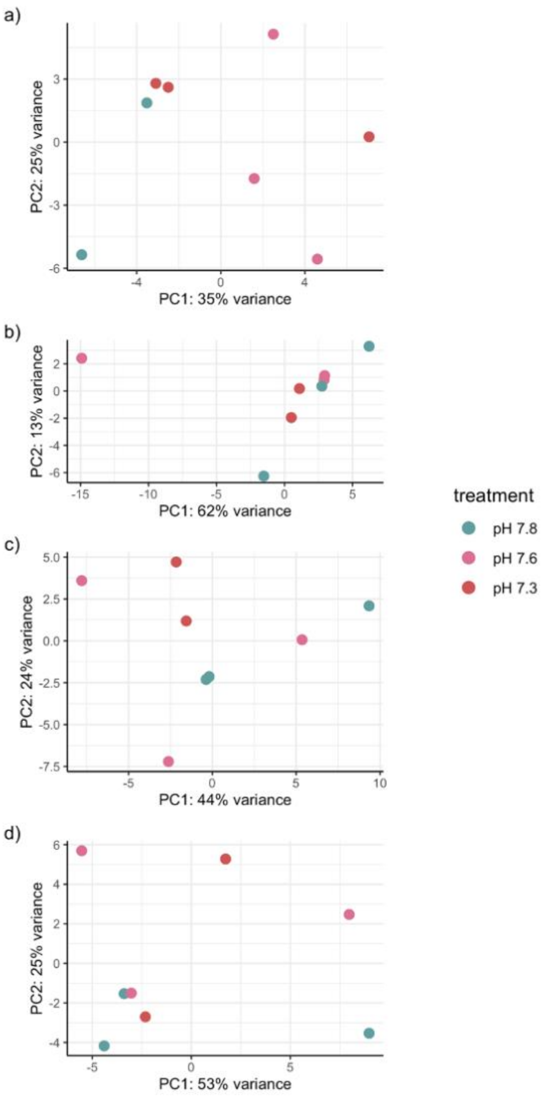

**Table S2.** Differential gene expression and k-means clustering (k=2) results for planula ([link](link))

**Figure S2.** Principal coordinates analysis of symbiont genes in the planula samples. a) A principal coordinates analysis based on sample-to-sample distance computed from the 1,365 genes passing a low counts filter, wherein a gene must have a count of 10 or greater in at least 7 out of 8 samples (pOverA ~0.875, 10). b) A principal coordinates analysis based on sample-to-sample distance computed from the 29 symbiont genes differentially expressed in planula developing in different pH environments.

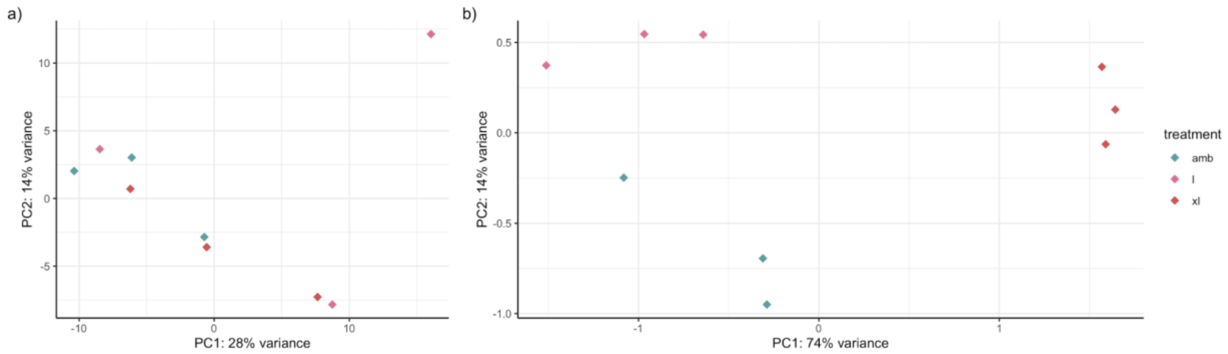


**Table S3.** Differential gene expression of symbionts extracted from planula holobiont mRNA samples ([link](link))

**Table S4.** Blastx alignment summary of the 29 differentially-expressed symbiont genes in the planula holobiont samples to the NCBI non-redundant database ([link](link))

**Table S5.** GOseq enrichment summary statistics with GOslim data and identifiers of associated differentially-expressed genes ([link](link))

**Table S6.** Kegg enrichment summary statistics and identifiers of associated differentially-expressed genes ([link](link))
